# MsPBRsP: Multi-scale Protein Binding Residues Prediction Using Language Model

**DOI:** 10.1101/2023.02.26.528265

**Authors:** Yuguang Li, Shuai Lu, Xiaofei Nan, Shoutao Zhang, Qinglei Zhou

## Abstract

Accurate prediction of protein binding residues (PBRs) from sequence is important for the understanding of cellular activity and helpful for the design of novel drug. However, experimental methods are time-consuming and expensive. In recent years, a lot of computational predictors based on machine learning and deep learning models are proposed to reduce such consumption. But those methods often use MSA tools such as PSI-BLAST or NetSurfP to generate some statistical features and enter them into predictive models as necessary supplementary input. The input generation process normally takes long time, and there is no standard to specify which and how many statistic results should be provided to a prediction model. In addition, prediction of PBRs relies on residue local context, but the most appropriate scale is undetermined. Most works pre-selected certain residue features as input and a scale size based on expertise for certain type of PBRs. In this study, we propose a general tool-free end-to-end framework that can be applied to all types of PBRs, **M**ulti-**s**cale **P**rotein **B**inding **R**esidue**s P**rediction using language model (MsPBRsP). We adopt a pre-trained language model ProtTrans to save the large consumption caused by MSA tools, and use protein sequence alone as input to our model. To ease scale size uncertainty, we construct multi-size windows in attention layer and multi-size kernels in convolutional layer. We test our framework on various benchmark datasets including PBRs from protein-protein, protein-nucleotide, protein-small ligand, heterodimer, homodimer and antibody-antigen interactions. Compared with existing state-of-the-art methods, MsPBRsP achieves superior performance with less running time and higher prediction rates on every PBRs prediction task. Specifically, we boost F1 score by 27.1% and AUPRC score by 7.6% on NSP448 dataset and decrease running time from over 10 minutes to under 0.1s on average. The source code and datasets are available at https://github.com/biolushuai/MsPBRsP-for-multiple-PBRs-prediction.

## 1. Introduction

Series of cellular functions, such as signal transduction, transport, immune reaction, metabolism, transcription and translation are driven by protein interactions [1]. Those interaction are carried out with a variety of other molecules including DNA, RNA, small ligand and other proteins [2]. Proteins interact with other molecules through a few binding residues on the interface [3]. Moreover, antibody-antigen interaction plays a crucial role in the immune response and is a specific protein-protein interaction [4]. Regions on the antibody and antigen that participate in the interaction are known as paratope and epitope, respectively [5]. Accurate detecting of protein interactions contributes to the construction of protein interaction networks [6, 7], annotation of protein functions [8], understanding of molecular mechanisms about diseases [9], design of new vaccines [10] and development of novel therapeutic antibodies [11].

With the success of the human genome project, millions of protein sequences are resolved and stored in public databases [12, 13]. However, the determination of protein interactions through traditional biological experiments is time-consuming and expensive [14–18]. Therefore, various computational methods have been proposed in recent years to solve this problem.

Based on the input data, computational methods can be divided into two groups: sequence-based and structure-based [1–3]. Just as the name implies, sequence-based methods only take protein sequence as input. And, structure-based methods usually utilize both protein sequence and structure [19–21].

Based on the level of prediction results, computational predictors can be categorized into two groups: protein-level and residue-level. The protein-level predictors judge a given pair of proteins interacting or not [22–25]. The residue-level predictors determine the positions of protein binding residues (PBRs). Sequence-based or structured-based computational methods can provide both protein-level or residue-level results.

Because there is a huge gap between the amount of protein sequences and structures. And protein structures of high resolution are not always available. Although structure-based computational methods can provide better results, they are also limited [26]. But sequence-based predictors are usually much cheaper and faster. Besides, residue-level results can provide more specific guidance for biological experiments. Therefore, we focus on the sequence-based computational methods predicting the positions of PBRs in this study.

According to recent reviews [1–3], there are various types of PBRs such as protein-protein binding residues, protein-nucleotide (DNA, RNA) binding residues, protein-small ligand binding residues and antibody-antigen binding residues. Table 1 summaries some methods predict different types of PBRs. We can find that all methods take advantage of machine learning-based or deep learning-based predictive models (shallow neural network, support vector machine, random forest, convolutional neural networks and recurrent neural networks, etc.).

**Table 1.**
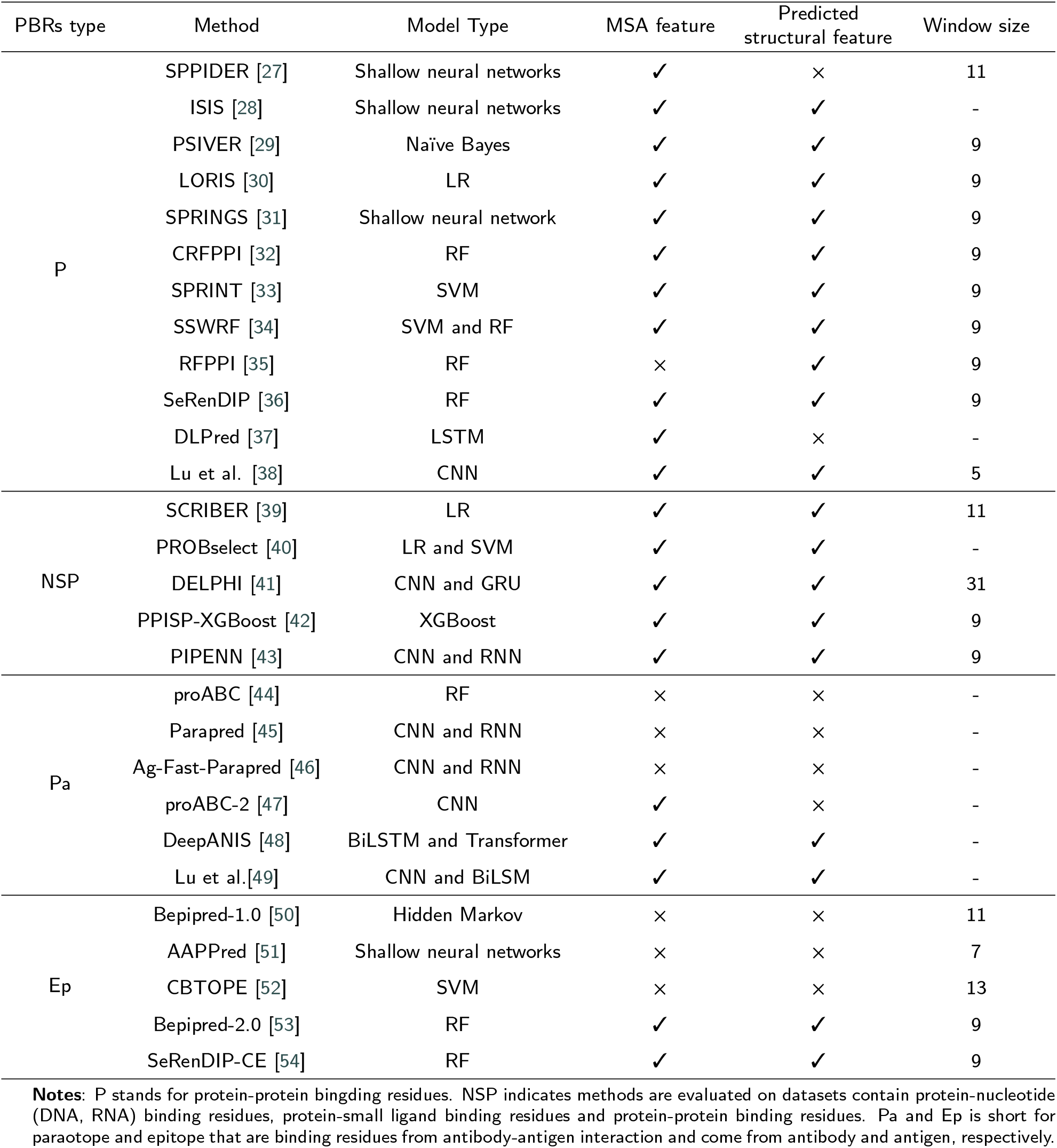
Summary of recent studies that focus on sequence-based PBRs prediction of different types.

From Table 1, we can observe that there are two main residue features including Multiple Sequence Alignment (MSA) feature and predicted structural feature that are used for input representation in most methods. MSA feature contains evolutionary information of protein sequence and can be obtained by running PSI-BLAST [55] or HHblits [56]. Compared to residue features driven from protein sequence, structural feature can provide richer information for input representation and make better prediction results. Therefore, a lot of sequence-based computational methods combine sequence feature with predicted structural feature such as secondary structure and solvent accessible surface by running NetSurfP-2.0 [57] or PSIPRED [58]. Some other common used residue features include physicochemical characteristics, physical properties, hydrophobicity and so on. The input of each protein in these methods is a matrix which is the concatenation of all residue feature vectors.

Meanwhile, the sliding window approach is popular in most methods for the aggregation of target residue and neighboring residues. This method simulates a window sliding over a protein sequence. And it concatenates a set of adjacent residue feature vectors representing the residue at center. It can be seen from Table 1 that different sliding window sizes are used for different predictors or predicting different types of PBRs. For example, 9 is the most common window size for predicting protein-protein binding residues. However, we test window size from 3 to 11 with step 2 in our previous work and achieve best performance with 5 [38].

In this paper, we introduce MsPBRsP, a general end-to-end sequence-based PBRs prediction framework without using any MSA tool. We can handle all types of PBRs, provide better prediction results and spend less running time comparing with the current predictors. There are three innovations in our work:

- We construct multi-scale features through using multi-size windows in attention layer, multi-size kernels in convolutional layer. This allows for better matching of different types of proteins and obtaining residue local context.
- MsPBRsP is capable of running on all types of PBRs and has been trained and tested on a number of benchmark datasets, one of them collects PBRs from protein interactions with nucleotide, small ligand and protein and is evaluated with a lot of PBRs predictors. The others collect PBRs from heteromeric protein, homomeric protein, antibody and antigen.
- To reduce the running time of input generation and provide comprehensive, uniform input data, we employ a pre-trained language model ProtTrans [59] on each protein sequence generating the input protein feature matrix. Running time is obviously reduced and results are significantly improved without running multiple sequence alignment and other structural feature predictors.

## 2. Materials and Methods

### 2.1. Benchmark datasets

To evaluate the performance of MsPBRsP on different types of PBRs and make fair comparison with the state-of-the-art methods [43], we collect various datasets from previous studies and utilize the same way for training and evaluation [36, 43, 45].

Table 2 shows the summary of all datasets used in this study. Among them, NSP6832 and NSP448 datasets contain multiple PBRs from protein interactions with nucleotide (DNA, RNA), small ligand and other protein. Other datasets consist of PBRs from type-specific interactions including protein-protein, heterodimer, homodimer and antibody-antigen.

**Table 2.** Summary of datasets.

| Dataset | Proteins | Residues | Binding Residues | PBRs type |
| --- | --- | --- | --- | --- |
| NSP6832 | 6832 | 2068793 | 221184 (10.69%) | NSP |
| NSP448 | 448 | 116500 | 22643 (19.44%) | NSP |
| P352 | 352 | 73188 | 11206 (15.31%) | P |
| P70 | 70 | 11791 | 2330 (19.76%) | P |
| He119 | 119 | 24328 | 3641 (14.97%) | He |
| He48 | 48 | 14056 | 1313 (9.34%) | He |
| Ho377 | 377 | 104914 | 22199 (21.16%) | Ho |
| Ho95 | 95 | 24766 | 5564 (22.47%) | Ho |
| Pa277 | 277 | 115341 | 6151 (5.33%) | Pa |
| Ep280 | 280 | 75956 | 6971 (9.18%) | Ep |

**Notes:** The meanings of P, NSP, Pa and Ep have been shown as notes in Table 1. He and Ho are short for heterodimer and homodimer, respectively.

The NSP6832 dataset is constructed for training the state-of-the-art method PIPENN [43]. The NSP448 dataset is generated by Zhang et al.[39] and widely used for testing and evaluating a lot of published predictors [1]. Protein sequences in NSP6832 and NSP448 interact with nucleotide (DNA, RNA), small ligands and proteins. And, a residue is defined as binding if the distance between an atom of this residue and any atom of its partner protein is shorter than 0.5 Å plus the sum of the Van der Waal’s radii of the two atoms. In this study, we utilize NSP6832 for training and NSP448 for testing and comparing with current predictors.

Although NSP6832 and NSP448 include PBRs covering most native protein binding residues [39], it should be noted that PBRs from type-specific protein interfaces may contain different properties. To evaluate performance on type-specific PBRs prediction, we utilize some other datasets containing specific protein interfaces.

P352 and P72 datasets are rebuilt from Dset186, Dset72 [29] and PDBset 164 [30] and are used for training DeepPPISP [20]. P352 and P72 contain common protein binding residues from heterodimeric interfaces. He119 and He48 include PBRs from homodimer interface and are used for training RFPPI_hetero [35]. It should be noted that He119 and He48 are subsets of Dset186, Dset72 [29]. Ho377 and Ho95 include PBRs from homodimer interfaces and are used for training RFPPI_homo [35]. And, residues are defined binding if both accessible surface area (ASA) before association and buried surface area (BSA) during the association are larger than 0 Å^2^ [35].

In a comprehensive review, antibody-antigen interface is regard as a specific type of protein interface [3]. Paratope and epitope are specific PBRs from antibody and antigen, respectively [5]. In recent years, a lot of computational methods for paratope and epitope prediction are proposed. Pa277 and Ep280 are popular benchmark datasets used for training and evaluating models [45, 54]. Liberis et al. [45] develop Pa277 which contains 277 complexes from Structural Antibody Database (SAbDab) [60]. Residues of antibody are defined as paratope if at least one atom is within 4.5 Å of any atom of the antigen. Ep280 is retrieved from SAbDab [60] as well by Hou et al. and includes 280 complexes [54]. A residue on antigen is assumed as epitope if any atom is less than 6.0 Å from any atom of the antibody.

The details of all datasets used in this study are present in the Supplementary File.

### 2.2. Input representation

To show the impact of different ways for residue features generation, we utilize two approaches to represent each input protein in this study. One is the usual way through residue features concatenation. The other way is similar to word embedding in natural language processing (NLP) and employing a pre-trained BERT model [61] named ProtTrans [59] which is trained on a super huge datasets Big Fantastic Database (BFD) [62, 63].

As shown in Fig.1A, we train MsPBRsP utilizing both input features (concatenated features and ProtTrans-based features). As shown in Table 1, MSA feature and predicted structural features are the most common used features. Therefore, concatenated features used in this study contains on-hot encoding of residue type, PSSM returned by running PSI-BLAST [55] and predicted structural features including secondary structure, solvent accessibility and backbone dihedral angles returned by running NetSurfP-2.0 [57]. Finally, each residue is represented as 52D vector which is the same as our previous work [38]. Because the superior performance in residue-level classification task [21] and lower running time than PSI-BLAST and NetSurfP-2.0, we employ pre-trained language model ProtTrans for feature generation. ProtTrans output a set of embedding vectors for each protein sequence. In this study, the input of MsPBRsP is a protein feature matrix *S* = {*r*_1_, *r*_2_, *r*_3_, …, *r*_*l*_},*r*_*i*_∈*R*^*d*^ (d=52 or 1024),where *l* is the protein length and *r*_*i*_ is the feature vector of i-th residue.

**Figure 1:**
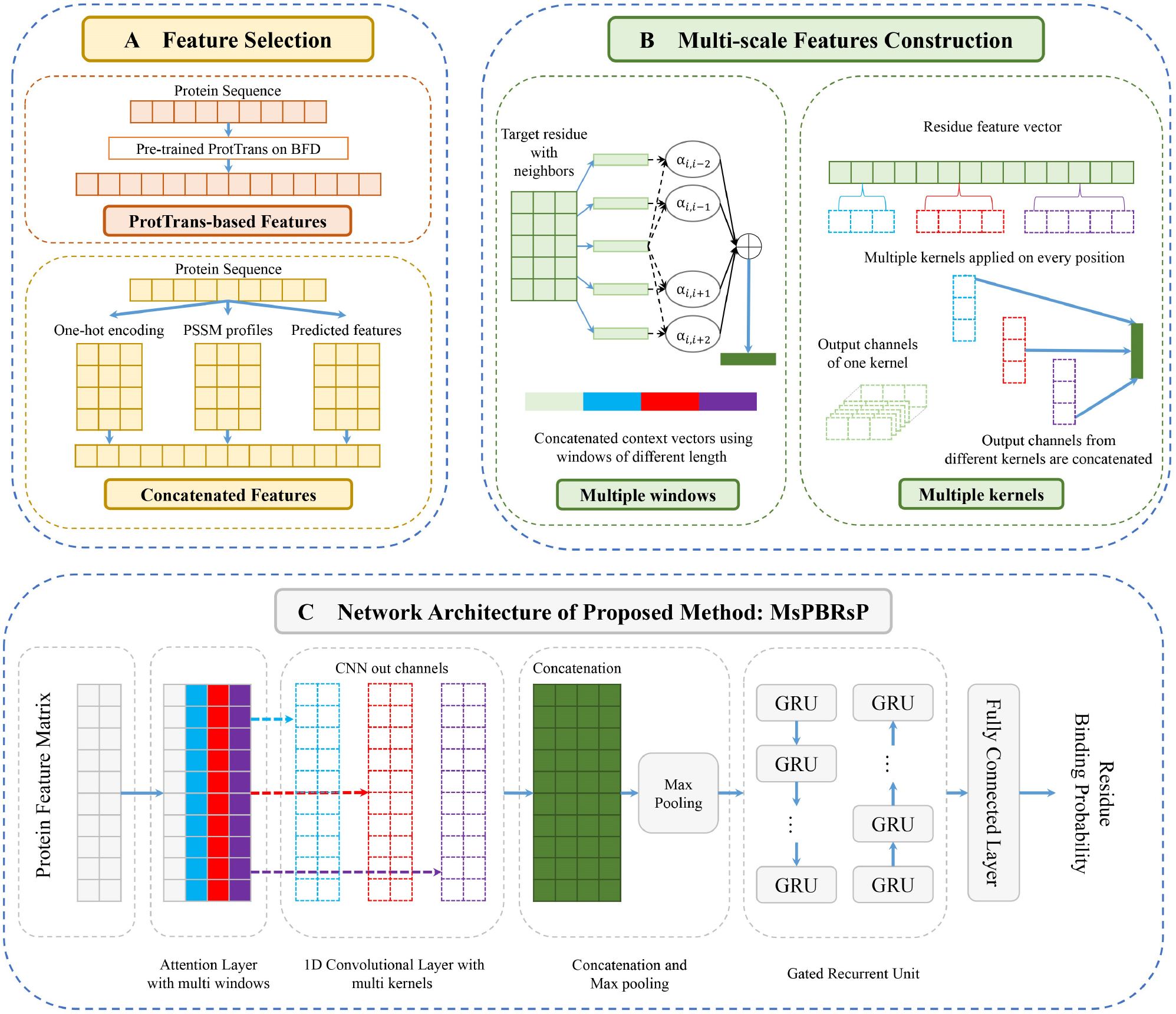
The framework of MsPBRsP. **(A)** Two ways of input feature generation used in this study for comparison and showing the effect of using pre-trained language model. **Upper**: ProtTrans-based features are returned through running ProtTrans which is trained on BFD. **Lower**: Concatenated features include one-hot encoding of residue type, PSSM profiles and predicted structural features such as secondary structure, solvent accessibility and backbone dihedral angles.**(B)** The construction of multi-scale features. **Left**: Context vector is generated by attention mechanism using sliding window approach. Multiple windows are used for adapting different types of PBRs and capturing complex patterns between target residue and neighboring residues. **Right**: Multiple kernels are utilized in convolutional neural networks on input residue feature vector for extracting richer information. The output channels of each kernel are concatenated. **(C)** Details of the framework architecture. Our proposed framework MsPBRsP mainly consists of attention layer, CNN layer and GRU layer.

### 2.3. Framework architecture

As shown in Fig.1C, MsPBRsP mainly contains three parts: an attention layer, a CNN layer and a GRU layer. The input is a protein feature matrix which is a set of residue feature vectors. The attention layer captures information from neighboring residues within multi-size windows. The CNN layer utilizes 1D convolutional operation with multi-size kernels and all output channels are concatenated and then max pooling operation is employed. The GRU layer contains bidirectional gated recurrent unit for extracting complex interaction relationship between all residues from the whole sequence. At last, a fully connected layer outputs the binding probability for each residue. Equations that specifically define MsPBRsP are listed in the Supplementary File.

#### 2.3.1. Attention Layer

The sliding window approach is useful for extracting information from neighboring residues in residue-level classification task [35]. However, the patterns of target residue and neighboring residue is complicated so that the window size for multiple PBRs prediction is different. Even for the same type of PBRs prediction, different best size may be used. The same conclusion can be reached from Table 1. Attention mechanism is helpful for aggregating target residues and neighboring residues which is proved in our previous work [38]. Moreover, the aggregation of target residue and neighboring performs best when the window size is 5 [38]. To adapt for complex properties of different type of PBRs, we construct multi-scale features by utilizing multi-size windows in attention layer.

As shown in the left part of Fig.1B, different weight is calculated for each neighboring residue and a context vector is constructed by weighted average of neighboring residues. And then, three window sizes (5, 15, 25) are used for capturing complex patterns between target residue and neighboring residues. The output of attention layer is the concatenation of original residue vector and three context vectors through using sliding windows of different sizes.

#### 2.3.2. CNN Layer

Convolutional neural networks (CNN) have been used in many bioinformatics tasks such as protein structure prediction [64], protein-protein interaction prediction [20] and protein-compound affinity prediction [65]. The usage of multi-size kernels in CNN is helpful for extracting information because it can account for various scales which has improved the performance in text classification [66] and image classification [67]. Inspired by TextCNN [66], we design a network architecture which expands on basic convolutional layer by using multi-size kernels in each 1D convolutional layer. This architecture allows the network to extract information over multiple scales. Therefore, we utilize three different kernel sizes (3, 5, 7) when carrying out our experiments. As Fig.1C shows, a concatenation operation is run resulting in feature matrix whose size is *L*×(3×1024) in which *L* is the length of input protein length and 1024 is the out channels of each 1D convolutional layer. And then, a max-pooling layer with filter size 5 and zero padding performs down sampling. The down sampled features are the input of the following GRU layer.

#### 2.3.3. GRU Layer

Recurrent neural networks (RNN) are very good at processing sequence data [68]. Both LSTM [69] and GRU [70] are variants of RNN and performance similarly, but GRU is computationally cheaper. The input of GRU layer is a set of residue feature vectors which has the same length with protein sequence. To capture complex interaction relationship among residues from the whole input protein sequence, we employ a GRU layer after the CNN layer. Similar with sentence in natural language, protein sequence also has a concept of forward and backward. Therefore, we utilize bidirectional gated recurrent unit. The size of the GRU’s hidden state was set to 64.

### 2.4. Implementation

We implement our proposed framework using PyTorch [71]. MsPBRsP is trained by minimizing the weighted cross-entropy loss using Adaptive Momentum (Adam) optimizer. The learning rate is tested among 0.1, 0.01 and 0.001. The training batch size is chosen from 32, 64 and 128. To achieve better performance and overcome overfitting, the dropout is tested among 0.2, 0.5 and 0.8. We utilize early stopping in the training process and performance drop on the validation set is detected [72]. For each combination, networks are trained until the performance on the validation set stops improving for 10 epochs. The parameters that maximize the area under the precision-recall curve (AUPRC) of the training data will be chosen.

Training time of each epoch varies roughly from 5 to 10 minutes depending on the sliding window length and the number the windows, using a single NVIDIA RTX2080 GPU. An independent validation set (10%of training set) is used to tune parameters. And 5-fold cross validation is employed when predicting paratope and epitope for making fair comparison with comparative methods.

### 2.5. Evaluation metrics

In this study, prediction performance is evaluated using eight classical binary classification metrics: Sensitivity (SEN), Accuracy (ACC), Precision (PRE), Recall (REC), Matthews Correlation Coefficient (MCC), F1, AUROC and AUPRC. Equations of all metrics are listed in the Supplementary File.

Because SEN, ACC, PRE, REC, MCC and F1 are all threshold-dependent metrics, we also use the area under receiver operating characteristics curve (AUROC) and area under precision-recall curve (AUPRC) which are threshold-independent metrics and provide evaluation on the overall performance of predictors. Moreover, AUPRC is more sensitive than AUROC when working with unbalanced datasets [73]. And the binding residues account for about 10% to 20% in the benchmark datasets used in this study. Therefore, AUPRC is the most import metric for framework evaluation and selection.

## 3. Results

### 3.1. Impact of multi-scales features

To extract richer information for improving framework performance on multiple PBRs prediction, we design multi-scale features through utilizing multi-size windows in attention layer, multi-size kernels in CNN layer and utilize GRU layer for capturing complex relationship from the whole protein sequence. Our proposed framework MsPBRsP utilizes 3 windows (5, 15, 25), 3 kernels (3, 5, 7) and 1-layer GRU. To evaluate impact of the number of windows, kernels and GRU layers, we employ experiments with the number of windows from 0 to 4, the number of kernels from 0 to 4 and GRU layers from 0 to 3. 0 indicates there is no attention layer, CNN layer or GRU layer. Architecture details of all methods mentioned in Table 3 are shown in Supplementary File. In this section, we train and test all models on NSP6832 dataset and NSP448 dataset five times to provide robust evaluation. The mean vale and standard error of all metrics are shown in Table 3.

**Table 3.** Impact of multi-scale features on NSP448 dataset.

|  | Methods | SEN | PRE | F1 | MCC | AUROC | AUPRC |
| --- | --- | --- | --- | --- | --- | --- | --- |
| Attention | W0K3GRU1 | 0.518(0.048)** | 0.636(0.010)* | 0.649(0.005)<sup>o</sup> | 0.297(0.011)<sup>o</sup> | 0.740(0.008)<sup>o</sup> | 0.425(0.007)<sup>o</sup> |
|  | W1K3GRU1 | 0.505(0.052)** | 0.641(0.007)* | 0.652(0.001)* | 0.303(0.001)** | 0.746(0.001)* | 0.428(0.002)* |
|  | W2K3GRU1 | 0.513(0.040)* | 0.641(0.007)* | 0.653(0.002)* | 0.305(0.005)<sup>o</sup> | 0.747(0.004)* | 0.429(0.002)* |
|  | W4K3GRU1 | 0.522(0.037)<sup>o</sup> | 0.630(0.010)* | 0.644(0.005)** | 0.288(0.011)** | 0.732(0.007)** | 0.410(0.009)** |
| CNN | W3K0GRU1 | 0.531(0.022)<sup>o</sup> | 0.632(0.001)** | 0.648(0.002)** | 0.295(0.003)** | 0.739(0.002)** | 0.413(0.004)** |
|  | W3K1GRU1 | 0.556(0.026)* | 0.632(0.003)** | 0.650(0.001)** | 0.299(0.002)** | 0.744(0.002)** | 0.419(0.002)** |
|  | W3K2GRU1 | 0.538(0.037)* | 0.637(0.004)<sup>o</sup> | 0.653(0.003)<sup>o</sup> | 0.305(0.006)* | 0.744(0.005)* | 0.428(0.006)<sup>o</sup> |
|  | W3K4GRU1 | 0.512(0.039)<sup>o</sup> | 0.639(0.003)** | 0.651(0.004)<sup>o</sup> | 0.302(0.008)<sup>o</sup> | 0.742(0.005)* | 0.423(0.007)* |
| GRU | W3K3GRU0 | 0.517(0.007)* | 0.637(0.007)<sup>o</sup> | 0.651(0.006)<sup>o</sup> | 0.301(0.013)** | 0.744(0.008)<sup>o</sup> | 0.428(0.008)* |
|  | W3K3GRU2 | 0.530(0.038)<sup>o</sup> | 0.627(0.002)** | 0.643(0.003)** | 0.285(0.005)** | 0.731(0.005)** | 0.405(0.004)** |
|  | W3K3GRU3 | 0.542(0.017)* | 0.616(0.002)** | 0.635(0.001)** | 0.268(0.003)** | 0.717(0.004)** | 0.380(0.007)** |
| Proposed | W3K3GRU1<br>(MsPBRsP) | **0.565(0.006)** | **0.645(0.005)** | **0.656(0.001)** | **0.311(0.002)** | **0.751(0.002)** | **0.433(0.001)** |

Attention: multi-scale features vary in attention layer. CNN: multi-scale features vary in CNN layer. GRU: multi-scale features vary in GRU layer. W indicates window. K indicates kernel. The number after W, K and GRU means the number of windows, kernels and GRU layers. The last row is our proposed method MsPBRsP. <sup>o</sup> denotes that the methods have no significant difference (two-sample t-test) with MsPBRsP. \*/\*\* denote that the methods are significantly worse than MsPBRsP with $P = 0.05/0.01$ , respectively. The best results are given in bold.

It can be seen from Table 3 that our proposed method MsPBRsP achieves best results on all metrics. Especially, there are significant decrease when no attention layer, CNN layer or GRU layer is used. The decrease declares the advantage of multi-scale features constructed in this study. Moreover, the prediction performances drop when there are 4 windows, 4 kernels or GRU more than 1 layer. The worse results indicates that the usage of multi-scale features should work for a certain framework depth.

The window size 5 is selected from our previous work for protein-protein interaction sites prediction [38]. Obvious difference is observed when window size is much larger than 5 [38]. Therefore, much larger window size 15 and 25 are used. It is obvious that 3 windows performs better than 1 or 2. The reason is that more useful information from neighboring residues at different distances are extracted. However, it may capture more distracted information when too larger window size is used such as 35 in W4K3GRU1.

Inspired by TextCNN [66], we utilize convolutional operation for extracting deeper features in MsPBRsP. However, information from adjacent residues has been extracted in attention layer. Therefore, we use 1D convolutional neural networks instead of 2D to capture information from residue-level feature vector. And the same kernel size is used in our proposed framework (3, 5 and 7). It can be seen from Table 3 that three kernels perform best and four kernels with size of 3, 5, 7 and 9 result in worse results. This situation is the same as the usage of multi-size windows. The cause is the introduction of more distracted information.

For capturing complex relationship from the whole protein sequence, we utilize GRU in MsPBRsP. As shown in Table 3, MsPBRsP with 1-layer GRU achieves better results than frameworks with 2-layer or 3-layer GRU. Moreover, the improvement is significant. This situation comes to the same conclusion with our previous work about structure-based epitope prediction [74]. Furthermore, linking GRU layer with CNN layer can improve the classification performance. The reason is that CNN-GRU framework includes both spatial and temporal layers. Therefore, convolutional perceptual representations and temporal dynamics simultaneously can be learned when CNN and GRU are jointly trained [67].

Finally, our proposed method MsPBRsP consists of three windows in attention layer, three kernels in CNN layer and 1-layer GRU. In summary, the usage of multi-scale features through using multi-size windows, multi-size kernels and GRU layer achieve best performance. Moreover, each component plays a import role for PBRs prediction.

### 3.2. Impact of ProtTrans-based features

The common way of input representation in a lot of bioinformatics tasks is concatenating a lot of residue feature vectors such as PSSM and predicted structural features. Because those features are returned by running PSI-BLAST [55] and NetSurfP-2.0 [57] which is time-consuming. And how many statistic results should be provided is not determined.To reduce the running time of feature generation and get comprehensive, uniform input data, we utilize ProtTrans-based features through employing a pre-trained language model on each protein sequence. To validate the advantage of the ProtTrans-based features, we employ MsPBRsP on concatenated features and ProtTrans-based features. And we train and test our framework on NSP6832 and NSP448 five times to provide robust results.

Fig.2 shows the comparison of concatenated features and ProtTrans-based features. From Fig.2A, we can see that it takes about 4000s to 5000s to run PSI-BLAST [55] and NetSurfP-2.0 [57] for various lengths of protein sequences from 100 to 700. If more residue features are used, it will take more time. However, the time of running ProtTrans [59] is hardly more than 0.1s which is significant faster. This obvious improvement indicates that we could utilize this feature on very large dataset, improve framework performance and make fast prediction.

**Figure 2:**
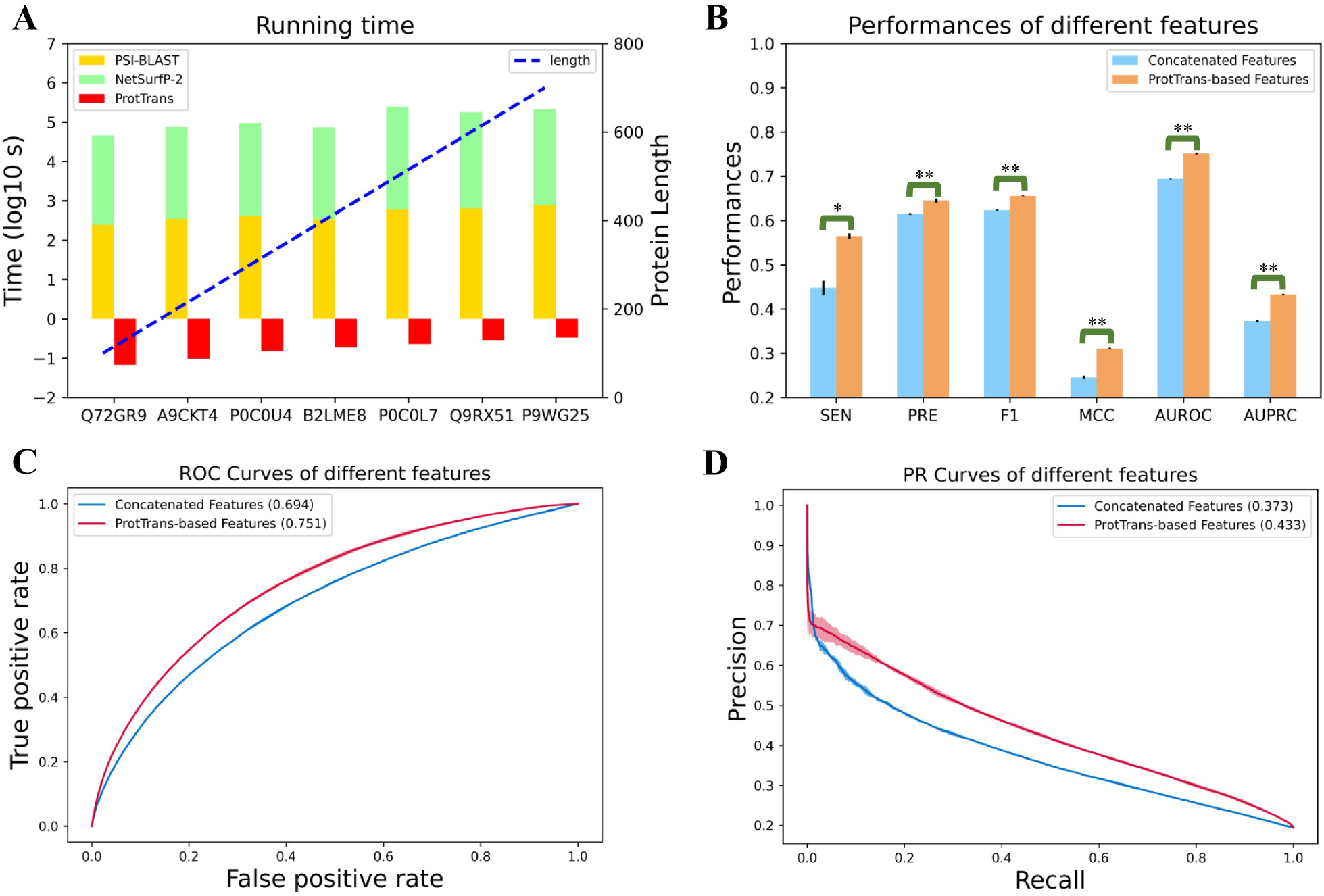
Comparison between concatenated features and ProtTrans-based features on NSP448 dataset. **(A)** The time of running PSI-BLAST [55], NetSurfP-2.0 [57] and ProtTrans [59] on various lengths of protein sequences. **(B)** Prediction performance comparison on SEN, PRE, F1, MCC, AUROC and AUPRC. We train and test MSPBRsP using both features five times. */** denote that concatenated features are significantly worse than ProtTrans-based features with P = 0.05/ 0.01. **(C)** The comparison of PR curves of two kinds of features. **(D)** The comparison of ROC curves of two kinds of features.

ProtTrans-based features also provide significant better results than concatenated features which shows the effect of transfer learning. Fig.2B shows average value and standard error of five experiments using two kinds of input features. It is obvious that ProtTrans-based features achieve significant better performance on all metrics. ROC cures and PR cures of all five experiments are shown in Fig.2C and 2D from which we can see that ProtTrans-based features improve 5.7%on AUROC and 6% on AUPRC than concatenated features.

### 3.3. Evaluation on NSP448 dataset

We compare the performance of MsPBRsP with other ten published sequence-based PBRs prediction methods on NSP448 dataset. MsPBRsP is trained and tested on the same datasets (NSP6832 and NSP448) with the state-of-the-art method PIPENN [43]. The results of all predictors are shown in Table 4 which are calculated on all residues from NSP448 dataset and called dataset-level performance in this study. The results of PIPENN [43] and DELPHI [41] are obtained from the published articles. And the results of other methods are collected from SCRIBER [39].

**Table 4.**
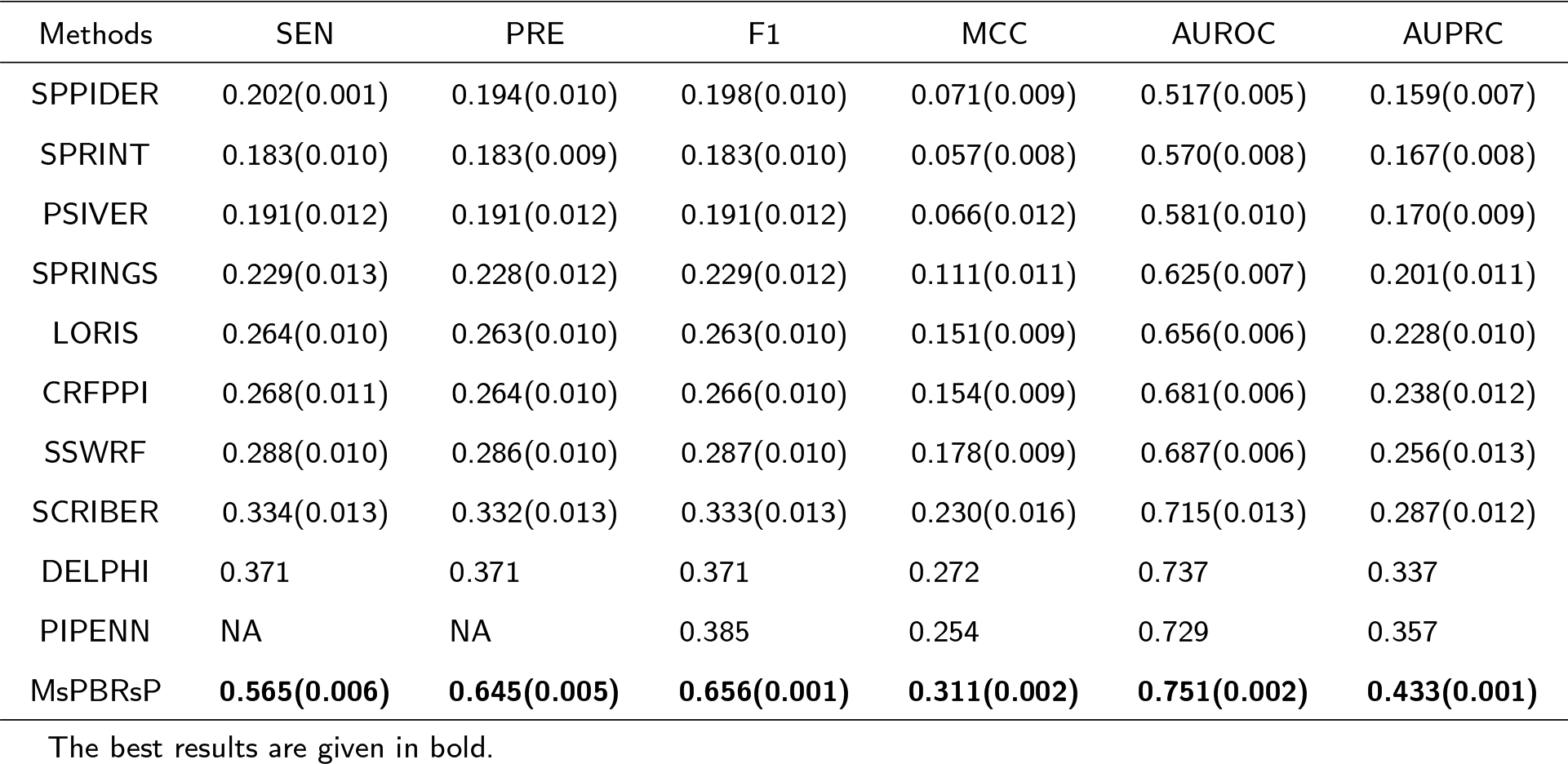
Comparison with competing methods on NSP448 dataset.

From Table 4 we can find that MsPBRsP achieves best results on all metrics. Especially, the improvement on the most important metric AUPRC is nearly 8% than the state-of-the-art method PIPENN [43]. F1 that evaluates overall performance also significantly improves, achieving 0.656 and there are 27.1% increase compared to PIPENN. MCC and AUROC of MsPBRsP reach 0.311 and 0.751 which corresponds to 3.9% and 1.4% improvement than DELPHI[41], respectively.

In order to make comparison with more predictors, Y. Li et al. [41] build NSP355 dataset by removing similar protein sequences in NSP448 dataset. Moreover, they provide the results of all comparative methods on each protein sequence in NSP355 at the website (https://delphi.csd.uwo.ca/). For making further comparison, we utilize the website service of PIPENN (https://www.ibi.vu.nl/programs/pipennwww/) gaining the prediction results of each protein sequence in NSP355. And then, we calculate the AUROC and AUPRC values of each protein. The protein-level results are shown in Fig.3 form which we can observe that MsPBRsP is the overall best predictor with median AUROC = 0.781 and median AUPRC = 0.504. The second-best method is PIPENN of which the median AUROC = 0.722 and median AUPRC = 0.372. And MsPBRsP achieves 5.9% and 13.2% improvement, respectively.

**Figure 3:**
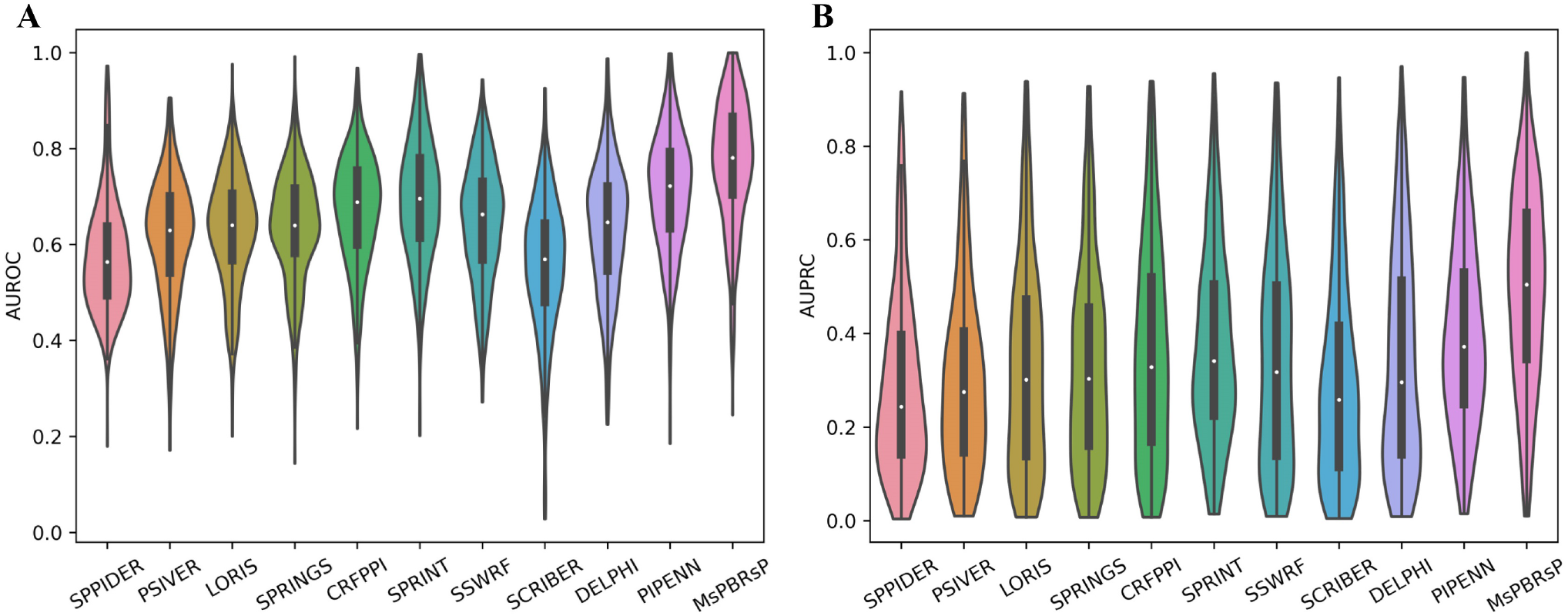
Protein-level performance comparison of MsPBRs and other methods on NSP355 dataset. **(A)** The distributions of per-protein AUROC values where the thick vertical lines represent the first quartile, median (white dot) and third quartile, whiskers denote the minimal and maximal values. **(B)** The distributions of per-protein AUPRC values as AUROC values.

### 3.4. Evaluation on type-specific datasets

As Table 5, 6 and Fig. 4 shows, we compare MsPBRsP with other methods on type-specific PBRs from protein-protein interaction, protein-nucleotide interaction, protein-small ligand interaction, heterodimer interface, homodimer interface and antibody-antigen interaction.

**Table 5.**
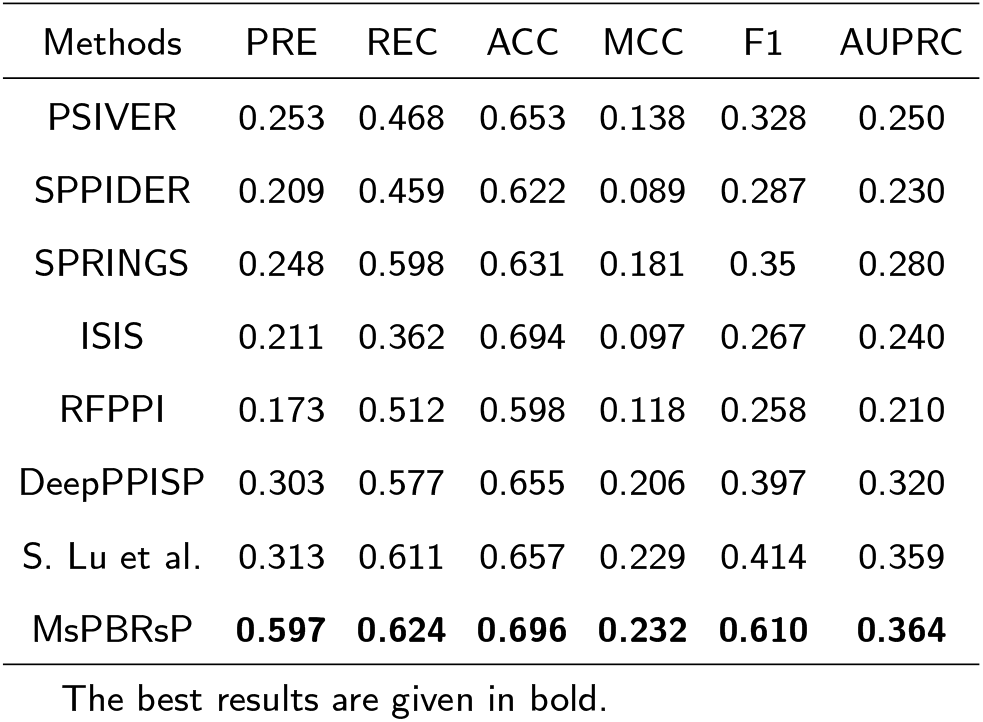
Comparison on P70 dataset.

**Table 6.** Comparison on type-specific datasets.

| PBRs Type | Methods | F1 | MCC | AUROC | AUPRC |
| --- | --- | --- | --- | --- | --- |
| S | SeRenDIP | 0.278 | 0.209 | 0.706 | 0.259 |
|  | PIPENN | 0.419 | 0.364 | 0.849 | 0.409 |
|  | MsPBRsP | **0.679** | **0.367** | **0.859** | **0.436** |
| N | DNApred | 0.294 | 0.198 | 0.609 | 0.240 |
|  | RNApred | 0.230 | 0.124 | 0.547 | 0.248 |
|  | PIPENN | 0.469 | 0.396 | 0.823 | 0.460 |
|  | MsPBRsP | **0.716** | **0.429** | **0.850** | **0.479** |
| He | SeRenDIP | NA | 0.122 | 0.636 | NA |
|  | PIPENN | 0.197 | 0.114 | 0.661 | 0.155 |
|  | MsPBRsP | **0.592** | **0.169** | **0.681** | **0.168** |
| Ho | SeRenDIP | NA | 0.277 | 0.724 | NA |
|  | PIPENN | 0.485 | 0.335 | 0.769 | 0.491 |
|  | MsPBRsP | **0.677** | **0.354** | **0.788** | **0.495** |

The best results are given in bold.

**Figure 4:**
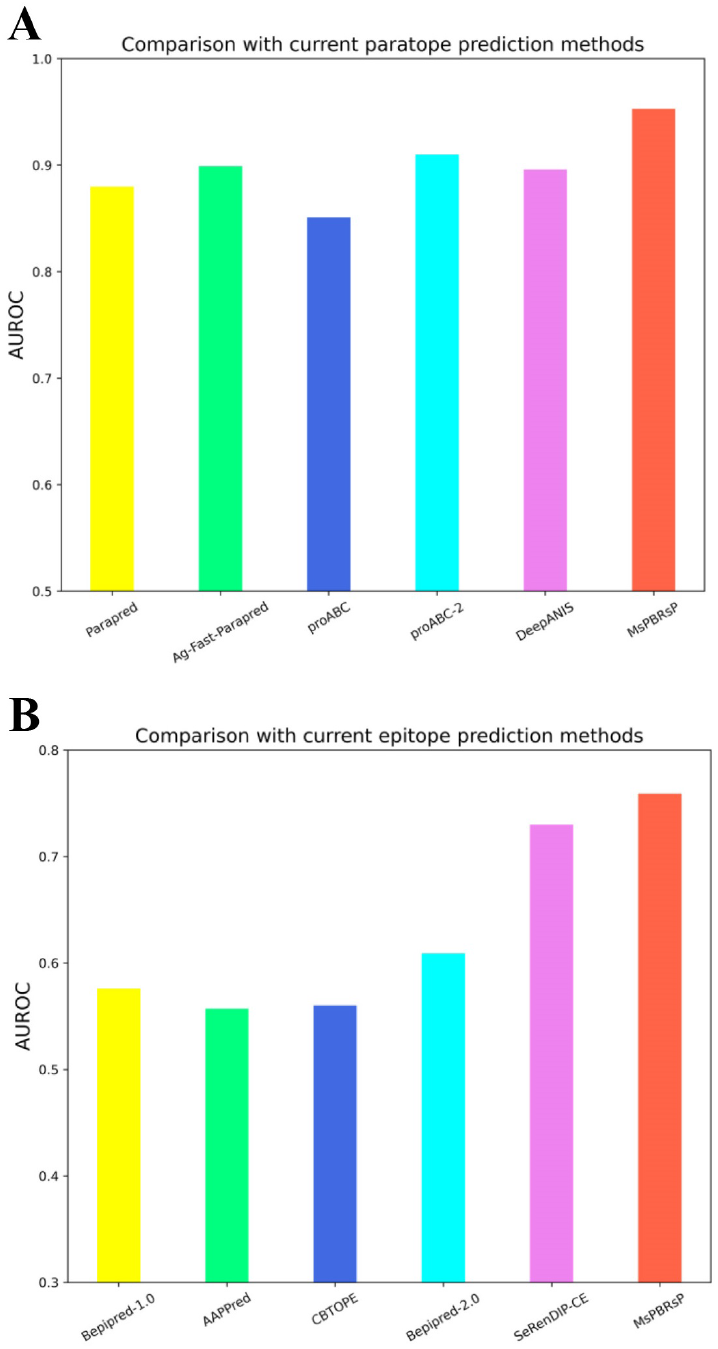
Comparison of MsPBRsP and other methods on paratope prediction and epitope prediction. **(A)** The AUROC values comparison of paratope prediction methods on Pa277 dataset. **(B)** The AUROC values comparison of epitope prediction methods on Ep280 dataset

To compare with more predictors, we train MsPBRsP on P352 and test on P70 which is the same way as DeepPPISP [20]. P352 and P70 are rebuilt from popular datasets Dset186, Dset72 [29] and PDBset 164 [30] which contain usual protein-protein interaction binding residues. The results of MsPBRsP and other competing methods are shown in Table 5. The performances of all other predictors are retrieved from DeePPISP [20]. It is obvious that MsPBRsP performs best on all metrics. MsPBRsB achieves 0.364 and obtains 4.4% improvement than DeepPPISP. Compared with our previous work which only utilizes one window in attention layer and mainly consists of attention layer and CNN layer, MsPBRsP performs significantly better on PRE, REC and F1.

In order to validate the advantage of PIPENN [43], B. Stringer et al. extract type-specific PBRs from NSP6832 and NSP448 for training and testing. The details of subsets from NSP6832 and NSP448 include PBRs from protein-nucleotide interaction and protein-small ligand interaction are shown in Supplementary File. In this study, we utilize the same way of training and testing MsPBRsP to predict type-specific PBRs. The results are shown in Table 6. Prediction results of comparative methods are retrieved from PIPENN. From Table 6, we can find that MsPBRsP achieves best performance on all metrics for predicting PBRs from protein-nucleotide interaction and protein-small ligand interaction. Especially, there are significant improvements of MsPBRsP than PIPENN on F1 which evaluates overall performance of predictor, correspond to 26% and 34.7% on protein-nucleotide binding residues and protein-small ligand binding residues.

Q. Hou et al. [35] build datasets including type-specific PBRs from heterodimer for training RFPPI_hetero. It should be noted that He119 is subset of Dset186 and He48 is subset of Dset72. Although we evaluate competing methods on rebuilt datasets of Dset186 and Dset72. For comparing with PIPENN [43], we still employ experiments on He119 and He48. Table 6 shows the comparison results from which we can find that MsPBRsP get highest values on all metrics. The comparison results further declare the ability of MsPBRsP to predict PBRs from heteromeric protein interaction. Ho377 and Ho95 are constructed by Q. Hou et al. [35] for training and testing RFPPI_homo which can detect PBRs from homodimer interface. We train and test MsPBRsP on the same datasets and the results are shown in Table 6. We can observe that MsPBRsP achieve best values on all metrics. The reason is that multi-scale features are useful for extracting different properties of PBRs from heteromeric proteins and homomeric proteins interaction.

Summaried by a comprehensive review [3], antibody-antigen interaction is an important and specific type of protein-protein interaction. Moreover, paratope and epitope are regarded as specific protein interface. To make further comparison and validate the advantage of multi-scale features used in MsPBRsP for handling multiple PBRs, we compare MsPBRsP with a variety of paratope and epitope prediction methods. All the comparative predictors employ multi-fold cross-validation on Pa277 or Ep280. In this section, we employ 5-fold cross-validation for training and evaluating MsPBRsP. The performance comparison on AUROC is shown in Fig.4. It can be observed that MsPBRsP performs best than all competing methods. The prediction results of comparative methods are retrieved from published works.

## 4. Conclusion

In recent years, various sequence-based PBRs predictors have been developed. However, there still are a lot of challenges for PBRs prediction based on protein sequence. For example, the MSA tools are needed to obtain protein residue information, which is time-consuming, and there is no unified standard definition of what statistics need to be obtained. Besides, the types of PBRs vary in the protein interaction objects such as nucleotide, small ligand, heteromeric protein, homomeric protein, antibody and antigen. It is difficult to determine the appropriate scale when extracting local residue information. Therefore, different types of PBRs contain different properties, which statistics should be selected when extracting residue features, and what scale of local residue information should be locked when predicting PBRs should be analyzed concretively. A more general and effective framework is needed to address these issues.

In our work, we propose a tool-free end-to-end deep-learning framework MsPBRsP which can provide general solution for multiple PBRs. Inspired by transfer learning, we employ a pre-trained language model ProtTrans [59] on each protein sequence in benchmark datasets for getting embedding features in this study. This will allow MsPBRsP to become a tool-free and end-to-end framework which makes prediction without running other tools and only takes raw protein sequence as input. The experimental results show that the usage of ProtTrans-based features in MsPBRsP significantly reduces running time of feature generation and improve prediction performance than concatenated features. To adapt for multiple PBRs, multi-scale features are designed and evaluated on various datasets collected in this study. Our proposed framework MsPBRsP mainly consists of attention layer, CNN layer and GRU layer. We construct multi-scale features through utilizing multi-size windows in attention layer, which is helpful to obtain the patterns of target residue and neighboring residue. The usage of multi-size kernels in CNN layer helps the network to extract more information. These benchmark datasets are collected from published studies and popular with a lot of PRBs predictors including protein-protein, protein-nucleotide, protein-small ligand, heterodimer, homodimer and antibody-antigen interactions. Lots of experiments are run on various benchmark datasets. The comparison results show that MsPBRsP outperforms a lot of comparative methods on predicting multiple PBRs. Specifically, on NSP448 dataset, F1 scores increased by 27.1% and AUPRC scores increased by 7.6%. On NSP355 dataset, AUROC increased by 5.9% and AUPRC increased13.2%.

Although MsPBRsP achieves best performances on multiple PBRs prediction, it has a limitation that partner information isn’t used. There are some proposed partner-specific PBRs prediction methods predicting binding residues for a particular pair interaction [75]. We will design novel deep learning model applying for partner-specific multiple PBRs prediction in the future.

## Acknowledgements

This work was funded by General Project of National Natural Science Foundation of China (61572444), National Key Scientific and Technological Project of National Health Commission of China (2019ZX09301-159), Bingtuan Science and Technology Project of Xinjiang Production and Construction Corps (2019AB034) and Leading Talents Fund in Science and Technology Innovation in Henan Province(194200510002)

