## Supplementary File for "MsPBRsP: Multi-scale Protein Binding Residues Prediction Using Language Model"

### Supplementary Materials

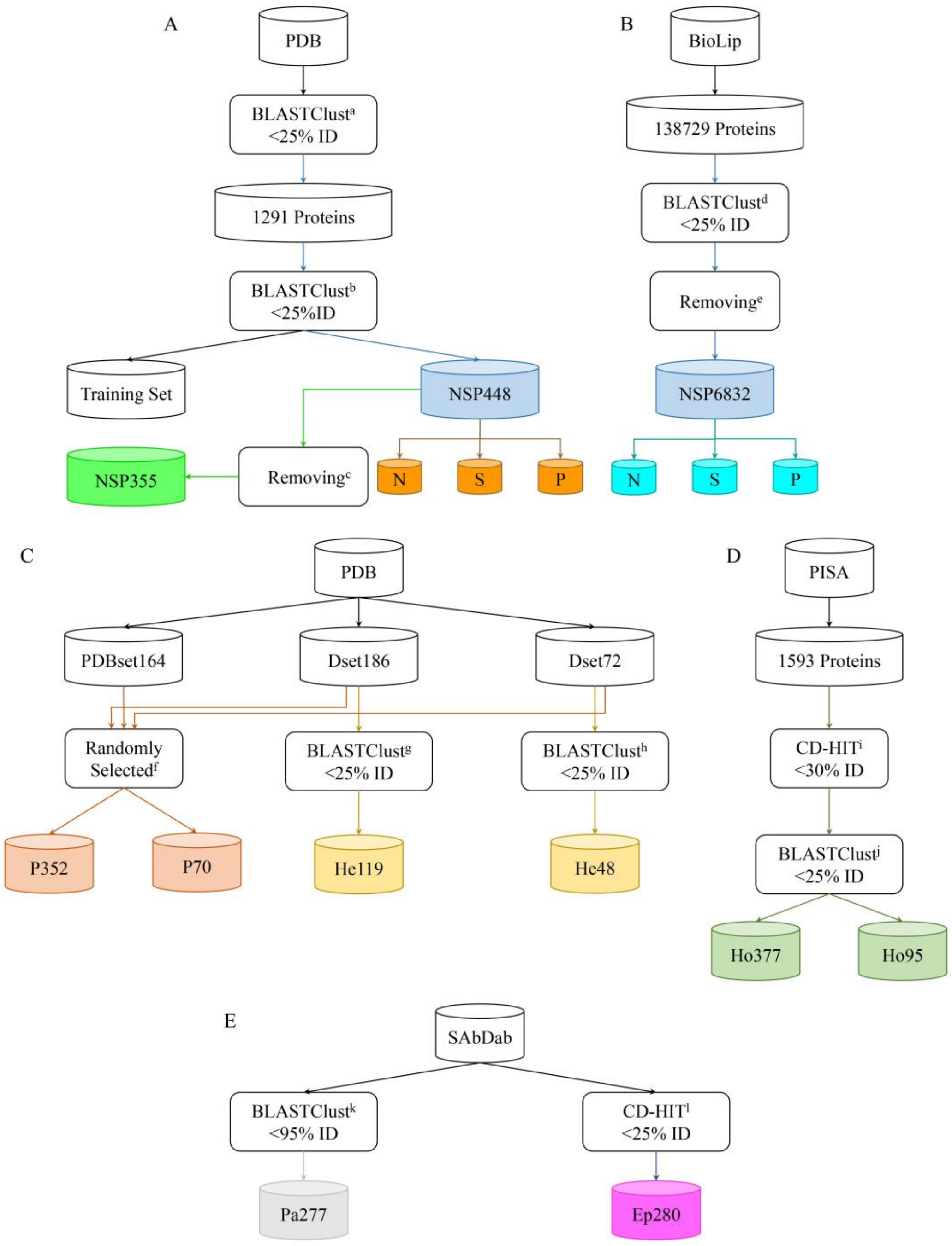


**Figure S1. Details of benchmark datasets used in this study.** (A): The steps to construct NSP448 dataset[1]. NSP355 is a subset of NSP448 which is constructed for competing with more methods[2]. NSP448 contains PBRs form protein-nucleotide (DNA, RNA) interaction (N), protein-small ligand interaction (S) and protein-protein interaction (P). The N, S and P subsets are used for evaluating performance for predicting type-specific PBRs. (B): The steps to construct NSP6832 dataset[3]. The N, S and P subsets of NSP6832 are used for training MsPBRsP as PIPENN to predict type-specific PBRs. (C): The steps to construct P352, P70[4], He119 and He48[5]

datasets which are used for training MsPBRsP to predict PBRs form heterodimeric interface. (D): The steps to construct Ho377 and Ho95 datasets[5] which are used for training MsPBRsP to predict PBRs form homodimer interface. (E): The steps to construct Pa277[6] and Ep280[7] which are used for training MsPBRsP to predict paratope and epitope.

**Notes:**

a: Protein sequences from UniProt are mapped to complexes from PDB and clustered to 1291 proteins by J.Zhang and L.Kurgan [8].

b: The remaining 1291 proteins are clustered with training sets from 7 competing models by J.Zhang and L.Kurgan [8].

c: Proteins with more than 40% similarity with DLPred’s training set[9] are removed by Y.Li et al.[2].

d: 138729 proteins from BioLip are clustered by B.Stringer et al. [3]

e: Proteins with the same UniProt Id in NSP448 and length shorter than 26 and longer than 700 are removed by B.Stringer et al. [3]

f: P352 and P70 are randomly selected from PDBset164, Dset186 and Dset72 by Zeng et al. [4]

g: Proteins in Dset186 are clustered with training sets of Dynamine and NetSurfP by Q.Hou et al. [5]

h: Proteins in Dset72 are clustered with training sets of Dynamine and NetSurfP by Q.Hou et al. [5]

i: 1593 proteins from PISA are filtered by Q.Hou et al. [5]

j: After filtered, proteins are clustered with proteins in Dset72 by Q.Hou et al. [5]

k: Clustered by E.Liberis et al. [6]

l: Clustered by Q.Hou et al. [7]

#### Evaluation Metrics

Evaluation metrics used in this study are shown as follows:

$$SEN = \frac{TP}{TP+FN} \quad (1)$$

$$ACC = \frac{TP+TN}{TP+FN+TN+FP} \quad (2)$$

$$PRE = \frac{TP}{TP+FP} \quad (3)$$

$$REC = \frac{TP}{TP+FN} \quad (4)$$

$$F1 = \frac{2*PRE*REC}{PRE+REC} \quad (5)$$

$$MCC = \frac{TP*TN-FP*FN}{\sqrt{(TP+FP)(TP+FN)(TN+FP)(TN+FN)}} \quad (6)$$

where TP (True Positive) or FN (False Negative) denotes the number of binding residues that are correctly or falsely predicted and FP (False Positive) or TN (True Negative) denotes the number of non-binding residues that are falsely or correctly predicted.

#### MsPBRsP Model

**Input**: The input feature matrix of each protein sequence, and $l$ is the protein length, $d = 52\ or\ 1024$ in this study.

$$S = [r_1, r_2, r_3, \dots, r_i \dots, r_l]^T, \ r_i \in R^d \quad (7)$$

**Attention Layer**:

The neighboring residue $r_j$ of target residue $r_i$ within window length are shown as:

$$r_j \in r_{i-w:i+w} = \{r_{i-w}, r_{i-w+1}, \cdots, r_{i-1}, r_i, r_{i+1,\cdots,} r_{i+w-1}, r_{i+w}\} \quad (8)$$

Multiple windows (5, 15, 25) of different length are used in MsPBRsP which are shown as:

$$r_j^1 \in r_{i-2:i+2} = \{r_{i-2}, r_{i-1}, r_i, r_{i+1,} r_{i+2}\} \quad (9)$$

$$r_j^2 \in r_{i-7:i+7} = \{r_{i-7}, r_{i-6}, \cdots, r_{i-1}, r_i, r_{i+1,\cdots,} r_{i+6}, r_{i+7}\} \quad (10)$$

$$r_j^3 \in r_{i-12:i+12} = \{r_{i-12}, r_{i-11}, \cdots, r_{i-1}, r_i, r_{i+1,\cdots,} r_{i+11}, r_{i+12}\} \quad (11)$$

The similarity score $score(r_i, r_j)$ between target residue $r_i$ and neighboring residue $r_j$, attention weight $\alpha_{i,j}$, the context vector $g_i$ of target residue $r_i$ are calculated as:

$$score(r_i, r_j)^n = V_a^n \tanh(W_a^n[r_i \oplus r_j^n ]) \quad (12)$$

$$\alpha_{i,j}^n = \frac{\exp(score(r_i, r_j^n))}{\sum_{j'} \exp(score(r_i, r_j^n))} \quad (13)$$

$$g_i^n = \sum_{j \neq i} \alpha_{i,j}^n r_j^n \quad (14)$$

$$r_i' = r_i \oplus g_i^1 \oplus g_i^2 \oplus g_i^3 \quad (15)$$

where n means n-th window.

**CNN Layer:**

The input of CNN layer is a concatenated matrix $S'$ shown as:

$$S' = [r_1', r_2', r_3', \cdots, r_{l-1}', r_l']^T, \quad r_i' \in R^{4d} \quad (16)$$

1D convolutional operation with multiple kernels are shown as:

$$c_i^{k^1} = f_c^{k^1}(W_c^{k^1} * r_i' + b_c^{k^1}) \quad (17)$$

$$c_i^{k^2} = f_c^{k^2}(W_c^{k^2} * r_i' + b_c^{k^2}) \quad (18)$$

$$c_i^{k^3} = f_c^{k^3}(W_c^{k^3} * r_i' + b_c^{k^3}) \quad (19)$$

where $k^1, k^2$, and $k^3$ mean different kernel sizes are used.

And then, max-pooling operation is applied on the concatenated out channels of each 1D convolutional layer:

$$p_i = maxpooling\ (c_i^{k^1} \oplus c_i^{k^2} \oplus c_i^{k^3}) \quad (20)$$

**GRU Layer:**

The out of CNN layer can be represented as a set of vectors which are shown as:

$$p_i = [e_1, e_2, \cdots, e_{3k}] \quad (21)$$

And the bidirectional GRU model can be show as the following equations:

$$z_t = \sigma(W_z \cdot [h_{t-1}, e_t] + b_z) \quad (22)$$

$$r_t = \sigma(W_r \cdot [h_{t-1}, e_t] + b_r) \quad (23)$$

$$\widetilde{h_t} = tanh(W_c \cdot [r_t * h_{t-1}, e_t] + b_c) \quad (24)$$

$$h_t = (1 - z_t) * h_{t-1} + z_t * \widetilde{h_t} \quad (25)$$

$$h_t' = \overrightarrow{h_t} \oplus \overleftarrow{h_t} \quad (25)$$

**Output**: The output of MsPBRsP is a fully connected network shown as:

$$o_i = f_o(W_o h_t' + b_o) \quad (26)$$

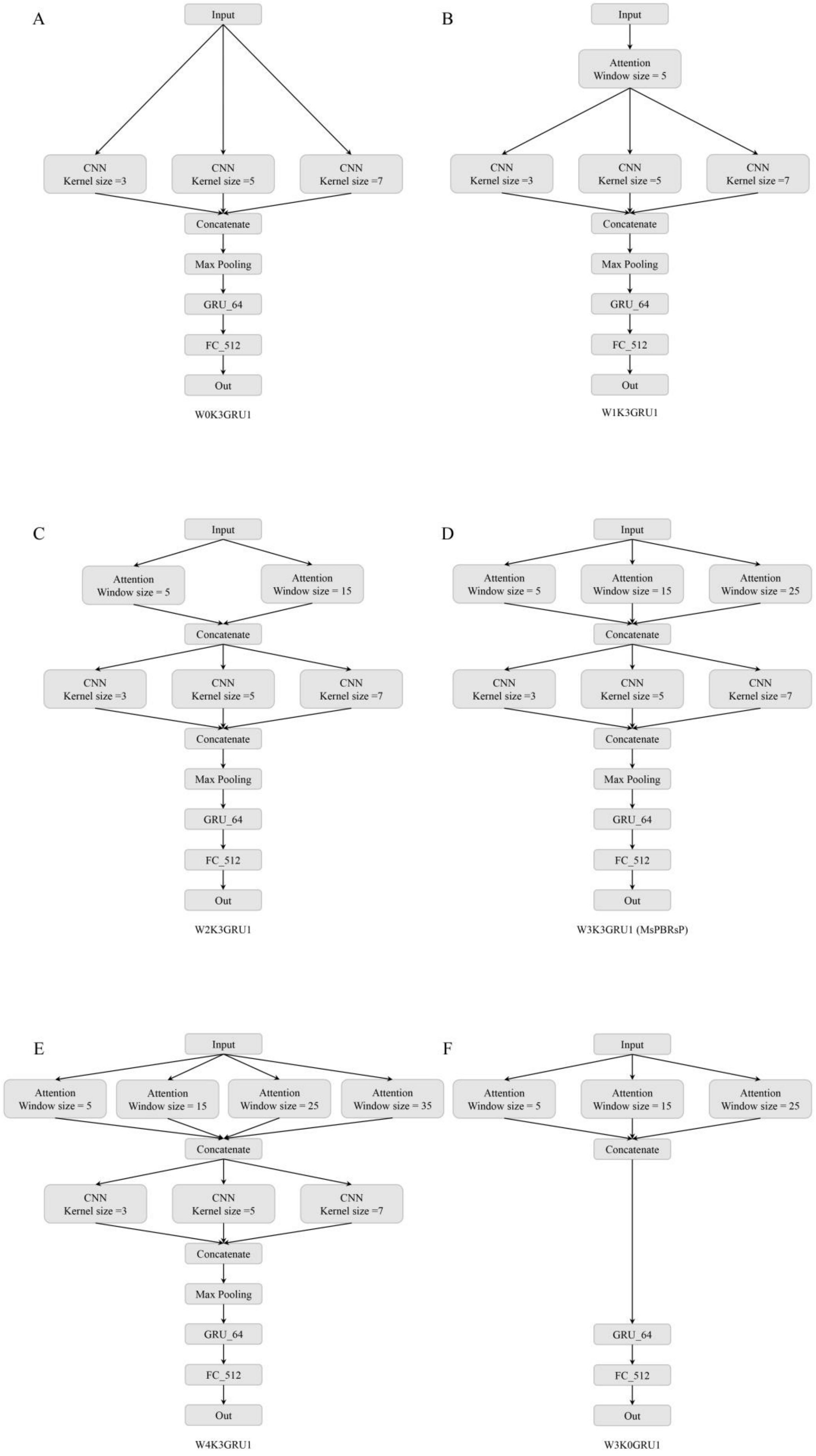


**Figure S2-1. Details of models mentioned in Section3.1.**

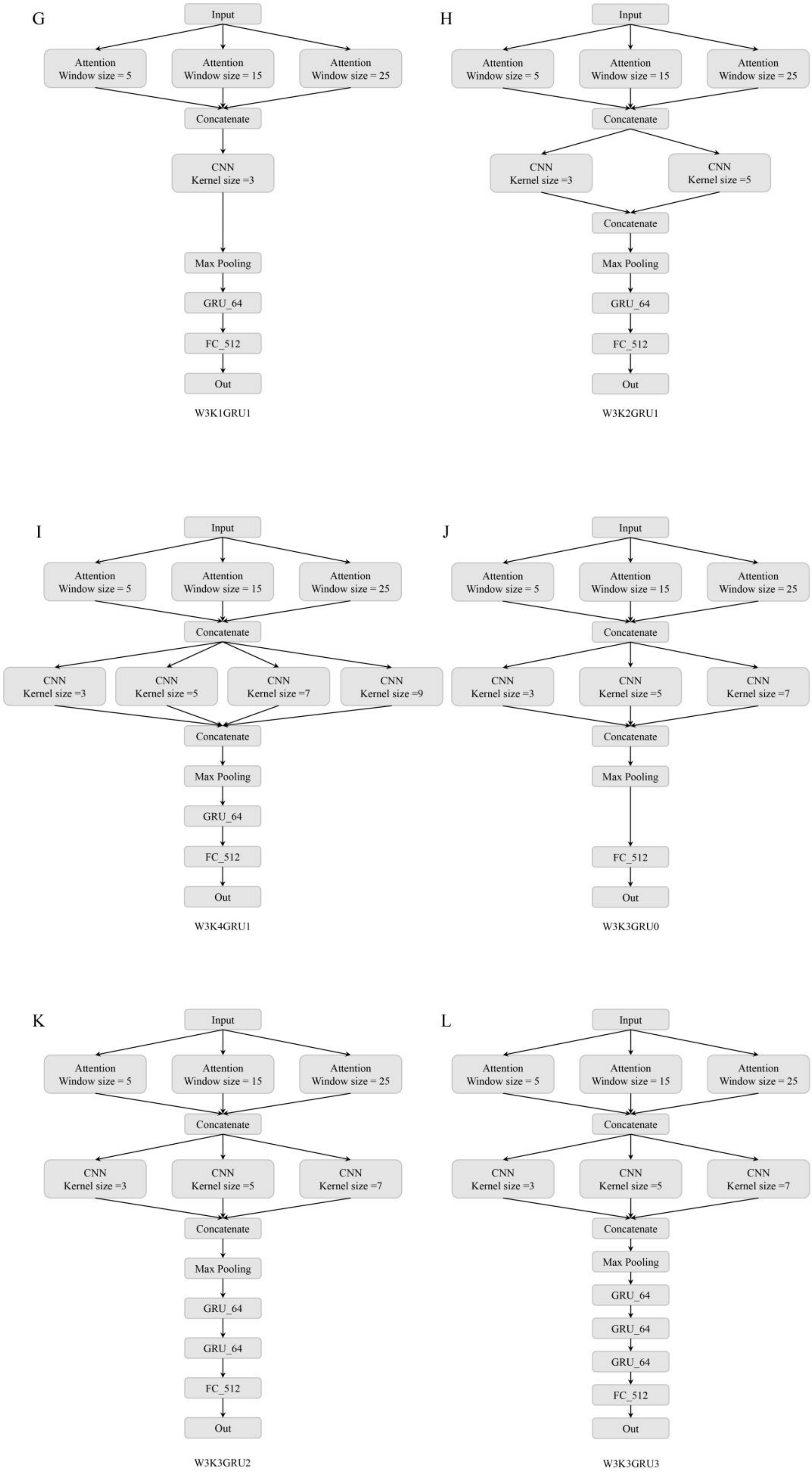


**Figure S2-2. Details of models mentioned in Section3.1.**

**Notes:** GRU_64: Bidirectional gated recurrent unit with 64 hidden nodes; FC_512: Full-connected layer with 512 hidden nodes.
